# Closely matched comparisons suggest that separable processes mediate contextual size illusions

**DOI:** 10.1101/2024.06.13.598943

**Authors:** Xinran A. Yu, Livia F. Fischer, Dietrich S. Schwarzkopf

## Abstract

Previous research suggests the magnitudes of the Ebbinghaus, Delboeuf, Ponzo, and tilt illusions all depend on the cortical distance between the neural representations of target stimuli and the surrounding context. However, several psychophysical studies found no compelling association between these illusions, calling this hypothesis into question. Here we ask if these discrepant reports could arise from methodological differences between these studies. We ran a battery of visual size illusion and basic discrimination tasks with carefully matched geometric properties, using a classic forced-choice design. In our small, homogenous sample, the Ebbinghaus and Delboeuf illusion magnitudes were strongly correlated, consistent with the idea that they reflect the same underlying mechanism when other sources of individual differences are minimised. Ponzo illusion magnitude also correlated with these two illusions, although less strongly in the case of the Ebbinghaus. Interestingly, the classic arrowhead version of the Mueller-Lyer illusion did not correlate with any of the other illusions or even with the objective ability to discriminate line length. This suggests that an altogether separate process underlies this perceptual effect. We further demonstrate that presenting stimuli briefly with central fixation critically affects measurements of the Ebbinghaus illusion. Additionally, we found that measuring illusion magnitude via adjustment is less reliable compared to two-alternative forced-choice procedures. Taken together, our findings suggest that different tasks probe separable processes determining illusion measurements. They further highlight the importance of the experimental design when testing relationships between perceptual effects and their links to neural processing.

## Introduction

The human visual system is not a digital camera. Our interpretation of visual space is subject to many contextual effects that can manifest as perceptual illusions. These effects can arise from basic limitations in neural architecture, such as the spatial resolution in peripheral vision being limited due to fewer neuronal resources than in central vision. In contrast, other illusions reflect the tasks the visual system has evolved to solve. For instance, navigating and interacting with a three-dimensional world involves interpreting the size of objects and their distance from the observer. Faced with a two-dimensional image that deliberately violates real-world relationships can lead to perceptual distortions.

### Mechanisms mediating contextual illusions

Several illusions of spatial context have been linked to the functional architecture and/or response properties of the visual cortex. Starting with a seminal study (Murray et al., 2006), numerous reports have established that activity in human primary visual cortex (V1) reflects the apparent, not the veridical, eccentricity of stimuli (Fang et al., 2008; He et al., 2015; Ni et al., 2014), the apparent size of retinal afterimages (Sperandio et al., 2012), the repulsive perceptual effect of adapting to different stimulus sizes (Pooresmaeili et al., 2013), and the perceived separation in a dot variant of the Mueller-Lyer illusion (Ho & Schwarzkopf, 2022). One study also suggested that effective connectivity between cortical representations of target and context stimuli correlates with the magnitude of the tilt illusion (Song, Schwarzkopf, Lutti, et al., 2013). In contrast to studies on the neural representation of illusions, we found that individual differences in the surface area of V1 correlate with the magnitude of the Ebbinghaus and Ponzo illusions (Schwarzkopf et al., 2011; Schwarzkopf & Rees, 2013) and the tilt illusion (Song, Schwarzkopf, & Rees, 2013). We found hints of a similar link also for the Delboeuf illusion (Moutsiana et al., 2016), although that study merely inferred illusion magnitude from raw perceptual biases, rather than measuring it via direct comparisons. Moreover, the relationship was significant only for some stimulus parameters. Taken together, these findings allude to a possible role of cortical magnification in mediating these contextual illusions: the cortical distance between target and context stimuli in V1 explains a substantial amount of the variance in the magnitude and direction of these visuospatial context illusions (Mareschal et al., 2010; Schwarzkopf & Rees, 2013; Urale & Schwarzkopf, 2023). Other indications that the Ebbinghaus illusion is at least partly mediated by early stages of visual processing are that it is substantially reduced when targets and inducers are presented to different eyes (Song et al., 2011) and it partly persists when inducers are masked from awareness by continuous flash suppression (Chen et al., 2018). These findings suggest a role for early, monocular channels.

The Ebbinghaus is often described as a “size contrast” effect whereby the perceived size of a target appears to change depending on the sizes of surrounding contextual stimuli (Massaro & Anderson, 1971). However, this interpretation is partly based on a naïve understanding of the effect in its classic configuration: a disc surrounded by a context of small discs (inducers), appears larger than the same disc surrounded by large inducers (Figure 1A). However, this interpretation misses the fact that when testing each inducer configuration separately against a no-context control, the illusion is in fact an underestimation of target size for most stimulus parameters (Roberts et al., 2005; Schwarzkopf & Rees, 2013; Urale & Schwarzkopf, 2023). Rather than size contrast, the illusion seems in large parts a geometric effect of the inducers, based on the interaction between contours (Todorović & Jovanović, 2018). The parameters exerting the most profound influence on the illusion are the distance between the target and the inducers, and how much of the surround is covered by inducers (Jaeger & Grasso, 1993; Jaeger & Klahs, 2015; Roberts et al., 2005; Urale & Schwarzkopf, 2023). This has led to the suggestion that the Delboeuf illusion is in fact a special case of the Ebbinghaus: an Ebbinghaus illusion with an annulus of very small, densely packed inducers is effectively a Delboeuf stimulus, yielding a nigh identical profile of illusion magnitudes (Roberts et al., 2005). According to this interpretation, results for these two illusions should be highly similar – that is, strongly correlated – across observers. Other studies have indeed found these two illusions to be highly similar, indicative of a role of contour attraction as the mediating mechanism (Sherman & Chouinard, 2016). The Ebbinghaus illusion is however more complex than the Delboeuf; the surrounding context consists of multiple stimuli of varying sizes, which are likely to interact with each other as well as the target. Moreover, there have been various suggestions of purportedly higher-level modulators of the Ebbinghaus illusion, such as the similarity between targets and inducers (Choplin & Medin, 1999; Deni & Brigner, 1997). While some of these effects are conceivably due to low-level feature interactions, certain studies suggest that semantic similarity may also play a role (Coren & Enns, 1993). The illusion effect likely involves both low-level contour interaction and higher-level processes related to size judgement (Jaeger & Klahs, 2015; Weintraub, 1979). If true, this implicates factors beyond local geometric interactions mediated by V1 circuits. This in turn predicts that a substantial proportion of the variance in the Ebbinghaus illusion is unrelated to that of the Delboeuf illusion.

**Figure 1.**
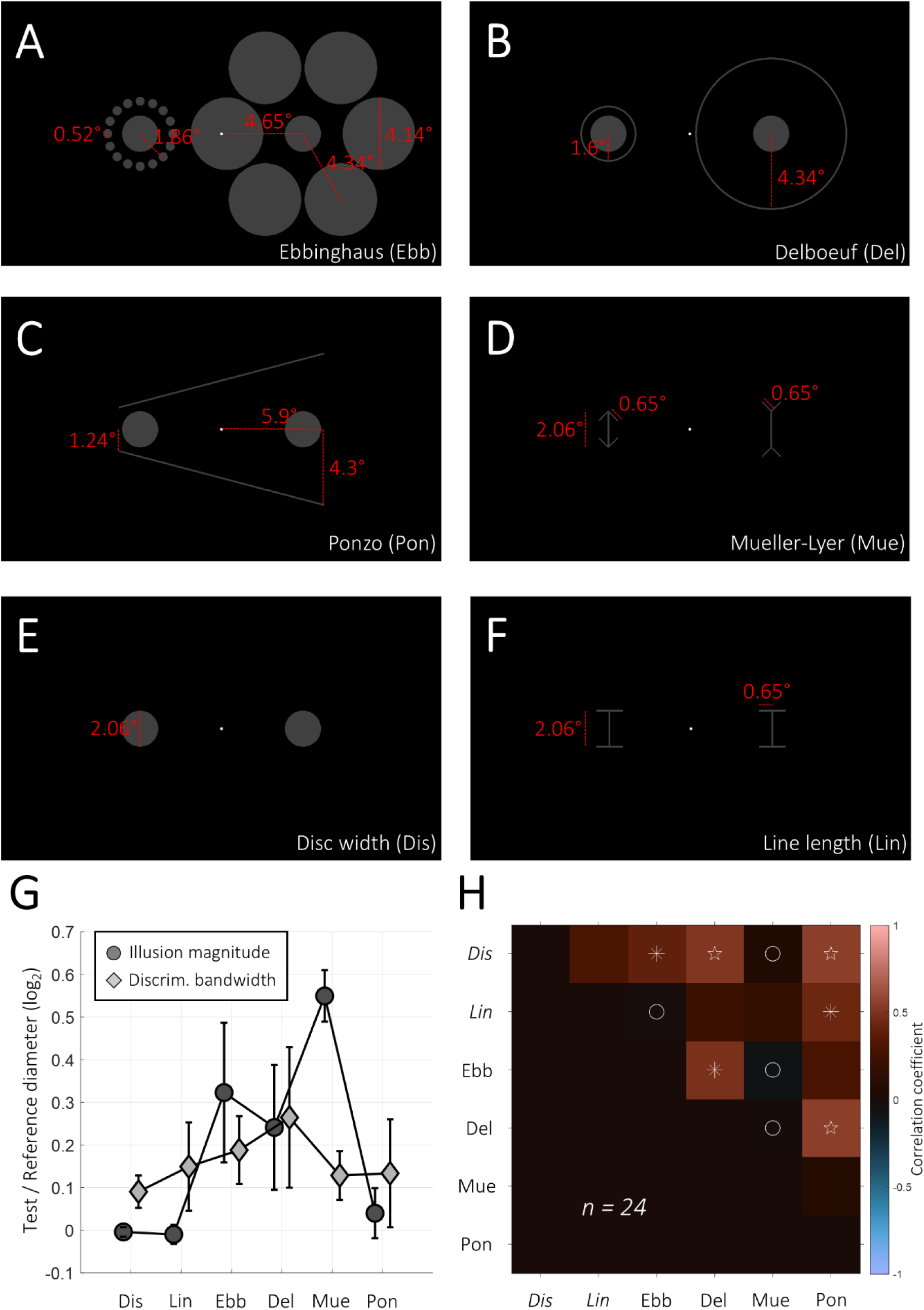
Stimuli (A-F) and results (G-H) of Experiment 1. In separate experimental runs, we estimated the illusion magnitude/perceptual bias and discrimination sensitivity for the Ebbinghaus (A), Delboeuf (B), Ponzo (C), Mueller-Lyer (D) illusions, or the discrimination of disc width (E) or line length (F). Red lines and numbers indicate dimensions in degrees of visual angle; these were not part of the presented stimuli. G. Average illusion magnitude (dark discs, solid lines) and discrimination bandwidths (light diamonds, solid lines) for each condition. Both measures are the ratio of the test relative to the reference stimulus diameter, expressed as binary logarithm (see Materials and Methods). Error bars denote ±1 standard deviation across observers. H. Correlation matrix comparing the magnitudes for the four illusions, Ebbinghaus (Ebb), Delboeuf (Del), Mueller-Lyer (Mue), Ponzo (Pon), and the discrimination bandwidths for judging disc width (Dis) or line length (Lin), respectively. Colours indicate strength of correlation. In each cell, symbols denote Bayes Factors: asterisk: BF_+0_>3, pentagram: BF_+0_>10, circles BF_+0_<⅓.

Similar questions arise about the Ponzo illusion (Figure 1C). Researchers often consider the Ponzo to reflect the interpretation of distance in depth from flat two-dimensional images, a failure of size constancy (Gregory, 1966, 2008). Such a process conceivably involves higher-order processing well beyond primary visual cortex, because it requires interpreting three-dimensional scenes and relies on neurons with larger receptive fields than those in V1. This hypothesis is borne out by findings that the Ponzo illusion exhibits complete interocular transfer (Song et al., 2011) and that it is wiped out when the context is rendered invisible through continuous flash suppression (Chen et al., 2018). Research, including our own, has also shown that both the classic Ponzo, a corridor-type illusion with real-world scenes, and various steps in between, all show an inversion effect: the illusion is substantially stronger if the context implies the ground rather than the ceiling (Altan et al., 2023; Brislin, 1974; Poom, 2020). One study also found that a version of the Ponzo illusion, in which the local geometry of the target remained constant, irrespective of whether it suggested a near or far interpretation, yielded the most pronounced illusion compared to other versions (Grzeczkowski et al., 2018). While there were still inevitable low-level differences between targets in this version, all these findings are better explained by a higher-level interpretation of the images than by a lower-level geometric interaction. While we found that the Ponzo illusion correlates with V1 surface area, just like the Ebbinghaus, this correlation was probably driven by an influential outlier and is therefore not robust (Schwarzkopf et al., 2011). Curiously, we also found no direct correlation between the Ebbinghaus and Ponzo illusion magnitudes themselves (but see e.g., Mazuz et al., 2023). Nevertheless, geometric parameters could theoretically play a role in the Ponzo. In the classic Ponzo configuration, the target which is typically perceived as smaller (and nearer to the observer) is generally positioned farther from the contextual stimuli; this conforms with the effect of target-inducer distance in the Ebbinghaus and Delboeuf illusions (Roberts et al., 2005; Urale & Schwarzkopf, 2023). Even in real-world corridor-type versions of the Ponzo, the distance between the “near/smaller” target and landmarks in the contextual image is greater. Interestingly, bandpass filtering the contextual images across different spatial frequency ranges does not impact the illusion’s magnitude (Yildiz et al., 2021). This suggests that while a purely low-level mechanism for this illusion is unlikely, it does not rule out a contribution of such processes.

Finally, in the classic Mueller-Lyer illusion, the lines are perceptually lengthened or shortened depending on the direction of arrowheads at the line ends (Figure 1D). This effect is often thought to involve higher-order analysis of three-dimensional scenes, such as interpreting the two configurations as edges/corners near or far from the observer (Gregory, 1966). However, evidence against this are the facts that the Brentano version of this illusion manifests as a perceived position shift and correspondingly the mid-point of the line, conflicting with the interpretation of lines as a cue for depth. The illusion is also effective for simple dot variants devoid of any lines (Ho & Schwarzkopf, 2022). The perception of this dot variant is reflected in V1 responses (Ho & Schwarzkopf, 2022), while the Brentano version is reduced when adapting to low, but not high, spatial frequency patterns (Carrasco et al., 1986). Nevertheless, the current literature cannot conclusively rule out a role of higher-order processing in this illusion, especially in the classic line-and-arrowhead version, which tends to produce stronger effects than stripped down versions like the dot variant (Coren, 1970). As with the Ponzo and the Ebbinghaus, it is likely that multiple separate mechanisms also contribute to the Mueller-Lyer.

Given this diversity, it seems unlikely to us that there could be a common factor to *all* spatial contextual illusions. Previous research searched for “common factors for visual illusions” (Cretenoud et al., 2021; Jastrzębowska et al., 2023), or specifically argued that “[…]*a person strongly susceptible to one visual illusion should also be strongly susceptible to other illusions*” (Grzeczkowski et al., 2017). These statements imply a factor that generalises across any visual illusions, even though the effects tested were conceptually rather diverse. Another study specifically compared a brightness contrast illusion to several diverse effects generally referred to as “geometrical illusions” (Axelrod et al., 2017). Interestingly, the magnitude of some of these illusions may be partly heritable (Coren & Porac, 1979) and related to personality traits (Makowski et al., 2023). Rather than variability in genuine perceptual appearance, this could indicate differences in how participants behave in psychophysical tasks, thus affecting the measurement. Another potential shared factor underlying such illusions could lie in visual *abilities,* like spatial acuity and sensitivity thresholds (Halpern et al., 1999), which could have downstream effects on illusion magnitudes (Coren & Porac, 1987). Yet, like for illusion magnitudes, correlations have been found only for few basic visual abilities (Cappe et al., 2014). A large psychophysical study on simultaneous contrast tasks found that the magnitudes of these illusions are mostly uncorrelated between stimulus dimensions (Bosten & Mollon, 2010), consistent with our own findings comparing size and luminance perception (Song, Schwarzkopf, & Rees, 2013). Often only versions of the same illusion (e.g., different configurations of the Ebbinghaus) share any significant variance at all, although such correlations are typically strong (Grzeczkowski et al., 2017). However, many of these reports used highly distinct stimulus parameters, making no effort to match conditions. Such an experimental design is reasonable for asking if there are any correlations between illusions in the broadest sense. For effects that arise in higher decision-making stages of cognition the exact stimulus parameters may be largely irrelevant. However, when testing hypotheses about the involvement of circuits in the early visual system, this is extremely problematic. If both the Delboeuf and Ebbinghaus illusion depend on the cortical distance between target and inducers, it is imperative that any comparison between these illusions controls for extraneous factors, rather than presenting each illusion at different visual field locations with different geometric parameters.

Experiments investigating these visual cortical processes therefore also require brief stimulus presentation and stable eye fixation. Both are essential for ensuring that stimuli are presented in fixed locations of the visual field and thus target the appropriate neuronal populations in retinotopic visual cortex. Yet, most studies exploring the relationship between different illusions used unconstrained experimental designs – again, a reasonable approach for studying the higher-order cognitive aspects of these tasks, but highly problematic for studying low-level perceptual processing.

### Measuring illusion magnitude

The ways how illusion magnitudes are measured is also critical. Any behavioural measures, including those quantifying perceptual experience, reflect the output of a whole brain working in concert. This ranges from multiple steps of image processing, making perceptual inferences, and eventually producing a motor output to report the judgement. This is another aspect in which no illusion is plausibly mediated solely by a single process, as any of these component processes can modulate the outcome. Numerous paradigms exist for quantifying the magnitude of an observer’s perceptual illusions, each of which involves cognitive confounds to some degree: this includes adjustment/reproduction tasks (e.g., Coren & Girgus, 1972; Ho & Schwarzkopf, 2022; Jastrzębowska et al., 2023), forced-choice procedures (e.g., Schwarzkopf & Rees, 2013), confidence ratings (Gallagher et al., 2019), and drift diffusion models (Sánchez-Fuenzalida et al., 2023). Adjustment tasks may be particularly prone to demand effects, because they rely on a subjective criterion for when there is a perceptual match between a test and reference stimulus, unlike forced-choice discriminations that require a directional judgement (e.g., which of two stimuli is larger). Reiterating an argument made by Morgan and colleagues (Morgan et al., 2013), one can imagine the difficulty of training a laboratory rat to perform an adjustment task. In contrast, the task is trivial to explain to human observers and may in fact be more natural and ecologically valid for how humans usually make perceptual judgements. And even simply by virtue of giving observers an extended time window to think carefully about their response, adjustment likely involves higher cognition than requiring a fast decision on briefly flashed stimuli.

However, it is now widely recognised that even classic forced-choice procedures – considered unbiased for measuring objective performance like acuity or detection thresholds (Green & Swets, 1966) – are susceptible to decision-making confounds. While classic forced-choice tasks allow the separation of the decision criterion from the discrimination sensitivity, when testing subjective effects like illusions such tasks cannot disentangle whether the decision criterion reflects perceptual or cognitive bias. This realization has led to the development of more complex procedures based on comparing the ‘difference between differences’ to disentangle perceptual from cognitive biases (Finlayson et al., 2017; Gallagher et al., 2019; Jogan & Stocker, 2014; Manning et al., 2017; Morgan et al., 2012; Morgan et al., 2013; Patten & Clifford, 2015).

Nevertheless, the classic two-alternative forced-choice (2AFC) design in which observers must make a perceptual judgement between a constant reference and a variable test stimulus remains a widely used method for quantifying illusion magnitudes. It constitutes a compromise between disentangling cognitive confounds for measuring perceptual bias and efficiency. It is however important that the observer’s expectations or preconceived notions of what the measurement reflects (such as suggestions that an illusion might be stronger/weaker in certain demographics) can be minimised. One way to achieve this is by randomly interleaving different conditions, randomising when/where reference and test stimuli are presented to control for response bias, and using a speeded design to prevent the observer from relying on non-perceptual cues in their decision. As explained, this design importantly also makes it relatively straightforward to control gaze and thus the low-level stimulus parameters.

We note that most of the studies that failed to find correlations between several contextual illusions used adjustment procedures (Cretenoud et al., 2021; Cretenoud, Francis, et al., 2020; Cretenoud, Grzeczkowski, et al., 2020; Grzeczkowski et al., 2017; Jastrzębowska et al., 2023). Adjustment tasks require prolonged stimulus presentation, during which observers tend to shift their gaze between stimuli while determining a perceptual match. Some previous experiments were also conducted online or outside lab environments, allowing minimal control over stimulus characteristics and eye gaze. Generally, adjustment tasks can be done in considerably less time than forced-choice paradigms. However, this confounds any conclusions we can draw from these findings.

For example, one of the studies that reported similar estimates for an adjustment and a 2AFC design (Cretenoud et al., 2021) found correlations between 0.6-0.67, meaning that between half to two thirds of the variance remained unexplained. The correlation in these experiments could largely reflect the higher-level decision processes but that the residuals – where the two measurements diverge – are due to the underlying stimulus-dependent processing by the visual system. While this is not explicitly stated in the methods, it is likely that these experiments did not control eye gaze and used unlimited stimulus presentations in the 2AFC tasks, further confounding any conclusions we can draw from these experiments about early visual processing. The illusion strength doubtless generalises across tasks up to a point, but the exact parameters – such as the retinal location of the stimuli – have a profound effect (Jaeger & Grasso, 1993; Jaeger & Klahs, 2015; Roberts et al., 2005; Urale & Schwarzkopf, 2023). Relatedly, a study on another geometrical illusion, the vertical-horizontal (bisection) illusion, demonstrated that gaze stability affects illusion strength (Chouinard et al., 2017).

### The present study

Here we set out to quantify the individual differences in these contextual size modulations, specifically the Ebbinghaus, Delboeuf, Ponzo, and Mueller-Lyer illusions. We tested a battery of these size illusions, using a classic two-alternative forced-choice (2AFC) procedure with brief stimulus presentations and fixation instructions. We also included two objective discrimination tasks for judging the width of discs and the length of lines, both without any illusion-inducing contexts. This was based on our previous research, which indicated associations between the objective acuities and subjective illusion magnitudes and associated spatial selectivity of visual neuronal populations (Moutsiana et al., 2016; Song et al., 2015; Song, Schwarzkopf, & Rees, 2013). The stimulus conditions across all these tasks were closely matched. For example, target stimuli were identical filled discs for testing the Ebbinghaus, Delboeuf, and Ponzo illusions as well as the corresponding disc-width discrimination tasks, and the length of the lines in the Mueller-Lyer illusion and the corresponding line-length discrimination task matched the diameters of the discs in the other tasks (Figure 1A-F). In the same vein, the distances between targets and inducers were kept comparable across the different illusions. This design therefore maximised the chance of finding associations between these perceptual effects; the influence of any extraneous stimulus differences should be minimised.

We tested a series of preregistered hypotheses aimed to better understand the mechanisms mediating these illusions. Critically, under the proposition that the Ebbinghaus and Delboeuf illusions are effectively the same phenomenon (Roberts et al., 2005), we predicted that these two effects are strongly correlated, directly contradicting another recent report (Jastrzębowska et al., 2023). Of note, a similar approach to ours – to minimise stimulus differences – also suggested a strong correlation between the Ebbinghaus and Delboeuf illusions when using an adjustment task (Sherman & Chouinard, 2016).

In a second experiment, we further assessed the effects of task and stimulus parameters on the Ebbinghaus illusion magnitude. We compared illusion magnitudes across three conditions: an adjustment task, a 2AFC task with unlimited stimulus presentation and free eye movements, and a classic 2AFC task with brief stimulus presentation and fixation instructions, as used in our earlier studies investigating brain-perception links for this illusion (Schwarzkopf & Rees, 2013). We also compared the effect of using two different stimulus parameters because the stimulus parameters described in our earlier study contained an error (see Materials and Methods for details), meaning researchers seeking to replicate those experiments following the methods would inadvertently use different stimuli than our original study (Jastrzębowska et al., 2023).

## Materials and Methods

Procedures were preregistered on the Open Science Framework for both Experiment 1 (https://osf.io/bhgud) and Experiment 2 (https://osf.io/5ey8h). While the study was concluded as preregistered, after several rounds of peer review, we agreed with the journal editor to double our sample size from 12 to 24. This additional data collection was also preregistered (https://osf.io/dmw4y).

### Participants

Twenty-seven observers (ages 18-45, 19 female, 2 left-handed) were recruited for this study, from both the staff and student body of the University of Auckland as well as the wider community. Three of these were the authors, while the remaining observers were naïve to the purpose of the experiments. Three observers were excluded from the whole study based on preregistered criteria (see below). Additionally, one observer was excluded from Experiment 2 only. Observers gave written informed consent, and all procedures were approved by the University of Auckland Human Participants Research Ethics Committee. Observers received a NZ$25 supermarket voucher for their participation in both experiments.

### Experiment 1

### Stimuli

Stimuli were closely matched across the different experimental conditions. In all tests, the fixation dot was a white disc (luminance: 140 cd/m^2^) with diameter of 0.15° visual angle, presented in the centre of a black screen (luminance: 0.6 cd/m^2^). All other stimuli were a dark grey (luminance: 6.2 cd/m^2^). The line width of any unfilled discs and line stimuli was 0.1°. All target stimuli were centred at an eccentricity of 4.65° to the left and right of fixation. The reference stimulus size (disc diameter or line length) was always 2.06°. The width or length of test stimuli varied on a binary logarithmic scale. The reference and test stimuli could appear either on the left or right of fixation. The main differences between the experimental conditions were in the design of the contextual stimuli (Figure 1A-F):

1. *Disc-width discrimination:* Only the target discs were presented. The stimulus design was identical to the control condition in our earlier study (Schwarzkopf & Rees, 2013), but note that the methods in that article incorrectly described the radius of the disc stimuli as diameters.
2. *Ebbinghaus illusion:* The target stimuli were filled discs, surrounded by other filled discs, the inducers, that were placed in equal angular steps, starting from the 3 o’clock position. Sixteen small inducers (0.52° diameter) were centred 1.86° from the centre of the reference stimulus. Six large inducers (4.14° diameter) were centred 4.34° from the centre of the test stimulus. These stimuli were also identical to the illusion stimuli in our earlier study (Schwarzkopf & Rees, 2013).
3. *Delboeuf illusion:* The target stimuli were filled discs, surrounded by rings. The ring around the reference and test stimuli had a radius of 1.6° and 4.34°, respectively.
4. *Ponzo illusion:* The target stimuli were filled discs. They were placed within a context of two converging lines that could either converge towards the left or right. The lines started and ended 5.9° horizontally from the vertical meridian in both directions. To converge, their ends were positioned above and below the horizontal meridian at a distance 4.3° and 1.24°, respectively. The reference and test stimuli were at the narrower and wider ends, respectively.
5. *Line-length discrimination:* The target stimuli were vertical lines. They bisected horizontal lines (1.3° long) at each end.
6. *Mueller-Lyer illusion:* The target stimuli were vertical lines. At each end, they were joined to two 0.65° long lines angled at ±45° relative to vertical, forming an arrowhead. In the test and reference stimuli, the lines were directed towards or away from the horizontal meridian, respectively.

### Procedure

After giving consent, participants were seated in front of a computer screen (Dell P2214H 21.5”;, dimensions: 47.6 cm x 26.8 cm, resolution: 1920 x 1080 pixels) at a viewing distance of 57 cm, with their heads stabilised by a chin and forehead rest. For the first set of participants (up until P12), the experiment was controlled by a Dell Latitude laptop 5410, using an integrated Intel UHD Graphics 770 card and running Psychtoolbox 3 (Brainard, 1997) in MATLAB R2023b under the 64-bit Windows 10 Education (version 22H2) operating system. For the remainder of data collection, we instead used a Dell desktop computer (Intel Cote i7-14700 2.1GHz CPU with 32GB memory) running Windows 11 Education (version 23H2). The general experimental design was a 2AFC task with an identical trial structure for all illusion and discrimination conditions. We matched stimulus parameters between the conditions as much as reasonably feasible. The task involved making a perceptual judgement about which target disc was larger or which line was longer, respectively. There were always two stimuli presented simultaneously within each trial, with one being a constant reference (diameter/length of discs/lines: 2.06°) and the other a test stimulus that varied across trials.

The experimental conditions were presented in the following sequence: participants were pseudo-randomised to start with either line judgements (line-length discrimination and Mueller-Lyer illusion) or the disc judgements (disc-width discrimination, and Ebbinghaus, Delboeuf, and Ponzo illusions) was. However, the objective discrimination tasks for these judgements always came first before testing the corresponding illusions. This familiarised observers – many of whom had little practice with visual psychophysics – with the task under simpler conditions before doing it under subjective modulations of size perception. The order of the three disc illusions was counterbalanced across participants, resulting in six possible permutations.

For each condition, the participant first performed a few demo trials in which the stimuli were presented until the participant’s keyboard response. This allowed the experimenter to explain the task. This was followed by more practice trials in which the trial sequence was as in the actual experiment, with 100 ms stimulus presentations. Once the participant confirmed they understood the task, the actual experiment began.

Each trial started with a brief 500 ms fixation period where only the fixation dot was presented on a black screen. This was followed by a 100 ms stimulus presentation, after which the screen turned completely black. The participant must then make a response to judge whether the left or right target stimulus was larger/longer by pressing the corresponding arrow key. The next trial then started immediately after the response.

The 160 trials were divided into four blocks of 40 trials each. There were four trial types: the two interleaved staircases (starting above or below true equality) and the reference stimulus could either appear to the left or the right of fixation. These four trial types were repeated 10 times per block in a pseudo-randomised order. Before each block, there was a rest screen reminding the participant of the task instructions and informing them of the number of blocks they had already completed. They commenced the block by pressing any key on the keyboard.

Stimulus levels were expressed in binary logarithms of the ratio between the test and reference size (diameter for discs, length for lines). This linearises stimulus values to account for Weber’s Law and allows the use of parametric statistical tests, which would not be applicable in the case of raw size ratios.

Test stimulus levels were controlled by two interleaved 1-down, 1-up staircase procedures that converged onto the point of subjective equality (PSE). One staircase started at a higher stimulus level (0.4 log units, wider disc or longer line) than true equality (0 log units), while the other staircase started at a lower level (-0.4 log units, narrower disc or shorter line). The staircase increased or decreased the stimulus level following the response in each trial. Initially, the increment/decrement was at a larger step size of 0.1 log units. Once five reversals had occurred in a given staircase, the step size was reduced to 0.05 log units. Moreover, from this point on, there was also a 10% probability on each trial that the stimulus level was increased or decreased (depending on the starting point of the given staircase) by 0.15 log units. These “;jitter trials”; prevented the stimulus level from continuously hovering around the PSE, as this could result in disengagement from the task.

### Data analysis

Observers completed 160 trials per experimental condition. Data were then analysed by calculating the proportion of trials in which the test stimulus was chosen at each stimulus level used (collapsing across both staircases in each condition and irrespective of which side the test was presented). We then fit a cumulative Gaussian function to these data points using the summed squared residuals approach, taking into account the robustness of each data point by weighting them by the number of trials at that stimulus level. We inferred the PSE from the stimulus level where the curve crossed the 50% response level (centroid of the psychometric curve). We also estimated discrimination uncertainty (conceptually a threshold or just noticeable difference) by determining the bandwidth of the psychometric curve.

Our preregistered plan was to exclude observers who failed to complete all experimental conditions; however, all observers completed all conditions. Further, we excluded participants for whom the bandwidth (technically, the σ parameter) of any psychometric curve fit exceeded 0.78 log units. Beyond this bandwidth, assuming the PSE is zero an observer would only score 10% or 90%, respectively, for stimulus levels of +/- 1 log units (i.e., test stimuli half or twice as large as the reference). We reasoned that this constitutes an implausibly poor performance for these simple visual discrimination tasks in observers with supposedly normal, healthy vision. All excluded participants were to be replaced to ensure correct counterbalancing of conditions (but see below for deviation from these criteria). Three observers were excluded based on this criterion.

The PSE (centroids of psychometric curves) for the illusion conditions and the discrimination abilities (bandwidths of psychometric curves) for the two objective discrimination tasks (disc width and line length) were compared with Pearson’s r correlation. We tested all hypotheses using directional (one-tailed) Bayesian correlation tests with custom MATALB code based on previously described procedures (Wagenmakers et al., 2016).

### Sampling plan

Our minimum sample size was set at N=6 because that is required to include all sequences for disc judgement illusions. We then incremented the sample size in batches of 6, accessing the stopping criterion after each batch. Using Bayesian hypothesis testing, this is an established procedure for sequential testing without introducing a statistical bias (Rouder, 2014). Our stopping criterion was based on hypothesis I_1_, that is, whether the one-tailed Pearson’s correlation between the Ebbinghaus and Delboeuf illusion magnitudes yielded a Bayes Factor of either 10 or 0.1, thus supporting either the alternative or null hypothesis, respectively. We chose this hypothesis for the stopping criterion because we expected these two illusions to reflect the same underlying neural process (Roberts et al., 2005; Schwarzkopf & Rees, 2013; Urale & Schwarzkopf, 2023), yet a previous study failed to find any strong correlation between them (Jastrzębowska et al., 2023). Our maximum sample size, in case the stopping criterion was not reached, was N=90. However, the criterion was reached after two batches were collected, that is, at N=12. This yielded conclusive evidence for many of our hypotheses. It is also only modestly smaller than the sample size used in many other similar lab-based studies comparing illusion magnitudes, which ranged from N=14-20 (Cretenoud et al., 2021; Grzeczkowski et al., 2017), N=30 (Jastrzębowska et al., 2023), to N=59 (Axelrod et al., 2017). Nevertheless, at the request of reviewers and the action editor, we collected an additional 12 participants, doubling our original sample size. The result of this study must therefore be considered partially exploratory; however, we preregistered this additional data collection. For transparency, we also present the results of the original preregistered sample in our public data repository (https://osf.io/5zs7y). Critically, the pattern of results remains qualitatively unchanged.

### Preregistered hypotheses

We tested the following hypotheses of between-subject correlations between psychometric parameters estimated for individual participants:

#### Hypothesis I_1_

Ebbinghaus and Delboeuf illusion magnitudes (PSEs) correlate positively, because they are effectively the same process.

#### Hypothesis I_2_

Ponzo illusion magnitude does not correlate (or only weakly) with magnitudes of the other illusions, because it reflects higher-level processing of three-dimensional scenes. (Note that this hypothesis contradicts exploratory hypothesis M3 below as we had no specific *a priori* predictions for the Mueller-Lyer illusion. If M_3_ had been confirmed, we would have expected I_2_ to be only partially confirmed)

#### Hypothesis S_1_

Ebbinghaus and Delboeuf illusion magnitudes each correlate positively with bandwidth for objective disc-width discrimination.

#### Hypothesis S_2_

Ponzo illusion magnitude does not correlate positively (or only weakly) with bandwidths for objective disc-width discrimination.

#### Hypothesis S_3_

Bandwidths for the objective discrimination tasks (disc width and line length) correlate positively because they both rely on local V1 processing (such as receptive field size).

#### Hypothesis M_1_

Mueller-Lyer illusion magnitude correlates positively with bandwidth for line-length discrimination.

#### Exploratory Hypothesis M_2_

Mueller-Lyer illusion magnitude correlates positively with the Ebbinghaus and Delboeuf illusion magnitudes if they share similar early neural mechanisms. Some previous findings suggest that this illusion could be mediated by local interactions in V_1_ (Carrasco et al., 1986; Ho & Schwarzkopf, 2022; Morgan & Glennerster, 1991), but this has not been tested using classic arrow-type Mueller-Lyer stimuli. Conversely, the correlation is only weak if this illusion reflects higher-order scene processing.

#### Exploratory Hypothesis M_3_

Mueller-Lyer illusion magnitude correlates positively with the Ponzo illusion, if they both reflect the same process of interpreting three-dimensional scenes.

A summary of the outcome of these hypotheses is shown in Table 1.

**Table 1.** Hypotheses tested in this study. *Ebb:* Ebbinghaus illusion magnitude. *Del:* Delboeuf illusion magnitude. *Pon:* Ponzo illusion magnitude. *Mue:* Mueller-Lyer illusion magnitude. *Dis:* Disc-width discrimination bandwidth. *Lin:* Line lentgh discrimination bandwidth. *Adjst:* Adjustment task. *Unco-2AFC:* Unconstrained-2AFC task.

| Experiment 1 – Comparison of several size illusions and related discrimination tasks |  |  |
| --- | --- | --- |
| Hypothesis |  | Outcome |
| I<sub>1</sub> | Ebb and Del correlate positively | **Supported** |
| I<sub>2</sub> | Pon does not correlate positively with Ebb, Del, or Mue | **Del: Rejected**<br>Ebb & Mue: Inconclusive |
| S<sub>1</sub> | Ebb and Del correlate positively with Dis | **Supported** |
| S<sub>2</sub> | Pon does not correlate positively with Dis | **Rejected** |
| S<sub>3</sub> | Dis and Lin correlate positively | Inconclusive |
| M<sub>1</sub> | Mue correlates positively with Lin | Inconclusive |
| M<sub>2</sub> | Mue correlates positively with Ebb and Del | **Rejected** |
| M<sub>3</sub> | Mue correlates positively with Pon | Inconclusive |
| Experiment 2 – Effect of task and stimulus conditions on Ebbinghaus illusion |  |  |
| Hypothesis |  | Outcome |
| H<sub>1A</sub> | Ebb differs between Fast-2AFC and Adjst | **Supported** |
| H<sub>1B</sub> | Ebb differs between Fast-2AFC and Unco-2AFC | **Rejected** |
| H<sub>2A</sub> | Ebb correlates positively between Adjst and Unco-2AFC | **Supported** |
| H<sub>2B</sub> | Ebb correlates less strongly between Adjst and Fast-2AFC | **Partially supported** |
| H<sub>3</sub> | Ebb differs between Grey&Large and White&Small | **Supported** but depends on task |

### Experiment 2

#### Stimuli

One of the Ebbinghaus stimulus designs (*Grey&Large*) was identical to that used in Experiment 1 and was a direct replication of our earlier study (Schwarzkopf & Rees, 2013). The other stimulus design (*White&Small*) was based on a replication study (Jastrzębowska et al., 2023), which was based on the incorrect method description in our original study (although we note that correct example stimuli were included in that study). Specifically, the spatial separation and eccentricities were identical to those in the other condition, but the filled disc stimuli were half width, because the methods description stated “diameter” where it should have said “radius.” For unexplained reasons, this replication study also presented stimuli in white instead of dark grey. Previous research suggests that white target stimuli are generally perceived as larger, which could impact the illusion estimate (Jaeger et al., 2014). Stimulus contrast also modulates contextual interactions in early visual cortex (Nauhaus et al., 2009), and could therefore affect the circuits we hypothesise mediate the Ebbinghaus illusion. To test this, we therefore also used white discs (luminance: 140 cd/m^2^) in this condition. Compared to the dark grey stimuli, many observers found these stimuli abrasive, especially when they were flashed briefly (see issues with reliability for this condition below). For Grey&Large disc stimuli, the fixation dot was a white dot as described in the preceding experiment; but for White&Small disc stimuli, the fixation dot was black as it fell inside an inducer.

#### Procedure

There were no demo or practice trials in this experiment, because participants had already done a very similar Ebbinghaus measurement in Experiment 1 immediately prior to this experiment. Instead, participants were shown images of the two possible stimulus parameters (Figure 2A-B) on the screen, and the experimenter explained again what the perceptual judgement entailed.

**Figure 2.**
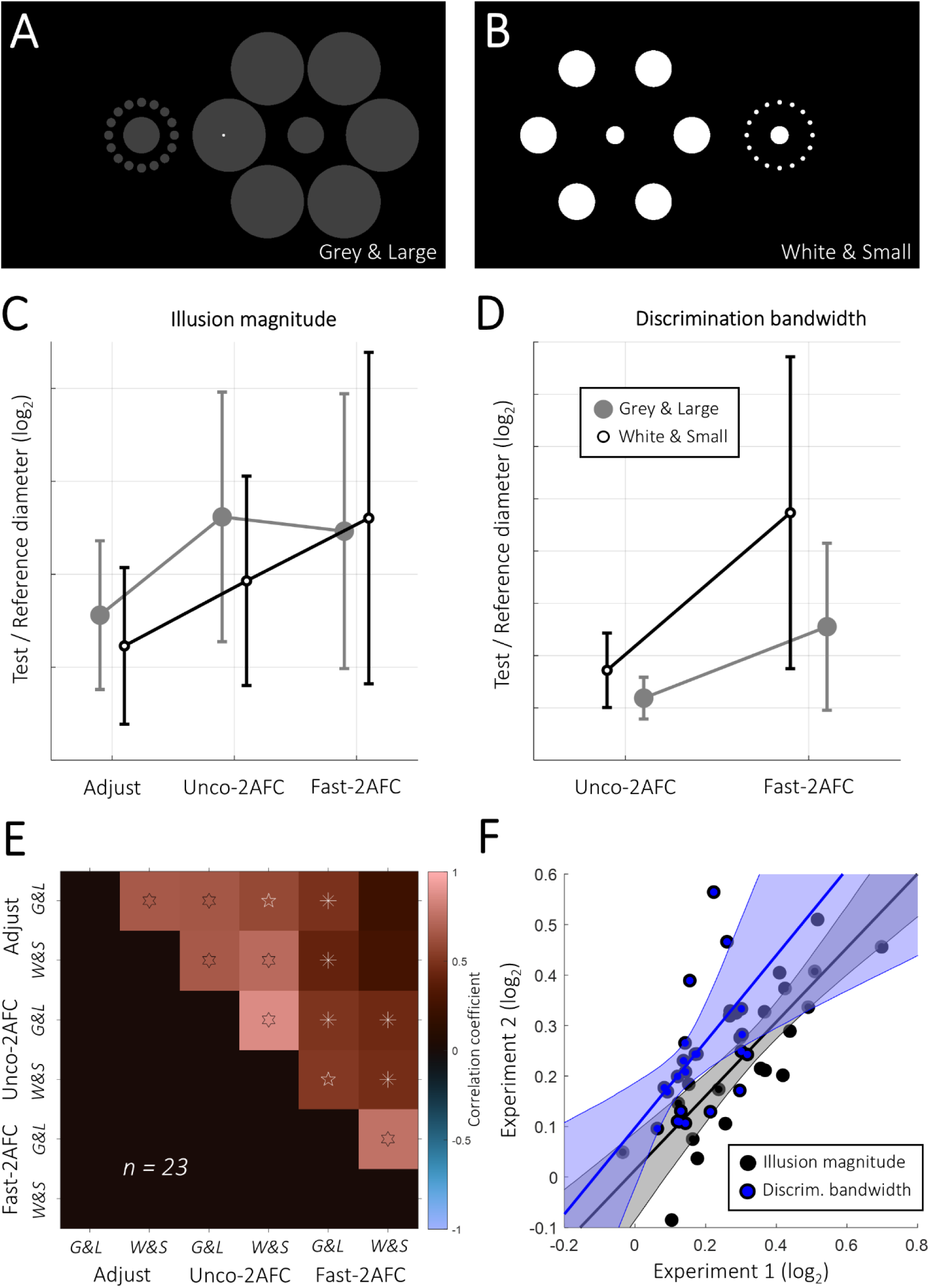
Stimuli (A-B) and results (D-F) of Experiment 2. We tested two configurations of the Ebbinghaus illusion: our standard stimulus from Experiment 1 (A) or another version with all discs half the diameter and in white (B). In separate experimental runs, we used an Adjustment, Unconstrained-2AFC, or a Fast-2AFC task to estimate the illusion magnitude and discrimination sensitivity (except for the Adjustment task). Average estimates for illusion magnitude (C) and discrimination bandwidths (D) for each task (see Materials and Methods and Figure 1 for details about these measures). Large grey discs denote the Grey&Large stimulus condition and small unfilled discs denote the White&Small condition. Error bars denote ±1 standard deviation across observers. E. Correlation matrix comparing the illusion magnitudes estimated for each condition (task and stimulus). Adjust: Adjustment task, Unco-2AFC: Unconstrained-2AFC task. G&L: Grey&Large, W&S: White&Small. Colours indicate strength of correlation. In each cell, symbols denote Bayes Factors: asterisk: BF_+0_>3, pentagram: BF_+0_>10, hexagram: BF_+0_>100. F. Illusion magnitude (black) and discrimination bandwidth (blue) estimated for the Fast-2AFC Grey&Large condition in Experiment 2 against the corresponding estimates in Experiment 1. Each dot denotes an individual observer. Solid lines show the linear regression fit and the shaded regions its bootstrapped 95% confidence interval using 10,000 iterations and the percentile method to derive confidence limits.

Participants then performed three tasks, with the order of tasks counterbalanced across participants, resulting in six permutations:

1. *Fast-2AFC:* This was essentially identical to the Ebbinghaus measurement in the preceding experiment, except for the inclusion of the White&Small disc condition. Participants were explicitly instructed to fixate on the white dot in the centre of the screen. There were 320 trials, divided into blocks of 80 (160 trials per stimulus condition, i.e. the same number as in the preceding experiment where there was only one condition).
2. *Unconstrained-2AFC:* This was the same as the Fast-2AFC task, but there was no fixation dot shown and the illusion stimuli remained on screen until the participant responded with the keyboard. They were explicitly told that they are free to move their eyes and inspect the stimuli as they please.
3. *Adjustment:* Stimuli appeared on the screen without a fixation dot. Participants then used the up and down arrow key on the keyboard to change the width of the test disc (surrounded by large inducers) until it was a perceptual match for the width of the reference disc. Once they were satisfied with the match, they pressed the space bar to move to the next trial. There were 20 trials in this test, 10 for each of the stimulus conditions. The side where the reference and test stimuli were presented was pseudo-randomised in each trial.

Stimulus values in the two 2AFC tasks were controlled by interleaved staircase procedures, just as in Experiment 1, except that there were independent staircases for the two stimulus conditions. In the Adjustment task, the initial radius of the test stimulus in each trial was drawn from a Gaussian distribution with a standard deviation of 0.4 log units.

### Data analysis

Data from the two 2AFC tasks were analysed as in Experiment 1. For the Adjustment task, the illusion magnitude in this experiment for each stimulus condition was calculated as the arithmetic mean across the final adjusted size of the test stimulus from all trials. The paradigm also collected information about the time course of the Adjustment task in each trial, but this is irrelevant for the present study and will not be discussed further.

Mean illusion magnitudes were compared using 2×2 repeated-measures analysis of variance (rmANOVA) using stimulus condition and task as factors. Post-hoc contrasts and paired two-tailed t-tests were used to compare individual conditions/tasks. The statistical evidence was based on default Bayes Factors. Correlations were calculated based on Pearson’s r, with statistical evidence assessed based on default Bayes Factors. Bayesian analysis of variance was conducted in JASP 0.18.3.0 (JASP Team, 2023), using a default Cauchy prior with scale r=√2/2 for comparisons of means. Bayesian correlation analysis was conducted with custom MATALB code using previously described procedures (Wagenmakers et al., 2016).

We also estimated the test-retest reliability of the illusion magnitudes in Experiment 2, using split-half correlations. For the Adjustment task, we calculated the illusion magnitude separately for odd and even numbered trials in each observer. For the 2AFC tasks, we refitted the psychometric curves separately for the two independent, pseudo-randomly interleaved staircases in each condition. We calculated the Pearson correlation, *r,* between the two halves across observers, and then extrapolated the reliability of the full dataset, *R_rel_,* using the Spearman-Brown prophecy formula (Spearman, 1910):

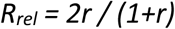

The magnitude of correlations between the Adjustment and 2AFC tasks was compared using the Williams/Steiger equation, which compares two correlations with a shared variable (Howell, 1997). Statistical inference was based on the 95% confidence limit of the Williams/Steiger t-statistic, derived by resampling pairs of observers 10,000 times with replacement. A difference in this directional test was deemed significant if the 95^th^ percentile of this bootstrap distribution was below zero. To compare correlations between stimulus conditions for the separate tasks, we bootstrapped the difference in correlation coefficients by resampling pairs of observers 10,000 times and calculating the 95% confidence level of this distribution. That is, this directional test was deemed significant if the 5^th^ percentile was above zero. We also used the same bootstrapping test to compare the reliability estimates for the different tasks.

Because we reused the same participants as in Experiment 1, the sampling plan for Experiment 2 was determined by the first experiment. We also applied the same exclusion criteria for Experiment 2, that is, if the bandwidth (technically, σ) of the psychometric curve fit for any of the experimental conditions exceeded 0.78 log units, observers would be excluded (without replacement, because recruitment was via Experiment 1). Based on this criterion, one observer was excluded, leaving only N=23 for Experiment 2.

### Preregistered hypotheses

#### Hypothesis H_1_

Illusion magnitude differs between the Adjustment task and the 2AFC tasks, specifically:

- *H_1A_:* Illusion magnitude differs between Fast-2AFC and Adjustment.
- *H_1B_:* Illusion magnitude differs between Fast-2AFC and Unconstrained-2AFC.

#### Hypothesis H_2_

Illusion magnitudes correlate between tasks as follows:

- *H_2A_:* Illusion magnitudes correlate between Adjustment and Unconstrained-2AFC.
- *H_2B_:* Illusion magnitudes correlate less strongly between Adjustment and Fast-2AFC.

#### Hypothesis H_3_

Overall, illusion magnitude differs between the stimulus parameters irrespective of task. Qualitatively similar effects are found comparing stimulus parameters in each task separately.

A summary of the outcome of these hypotheses is shown in Table 1.

### Deviations from preregistration

Our preregistered protocol also included a fourth hypothesis about the correlations between stimulus parameters. However, after data collection had commenced, we noticed that one sub-hypothesis (H_4B_) in the preregistration was phrased incorrectly, effectively repeating hypothesis H_2B_. The methods also did not include a description of the statistical comparison required for these independent correlations. Even though this was intended to be a preplanned analysis, we therefore decided to label this as exploratory to avoid the impression of altering the preregistration after the results were known.

We also adjusted our exclusion criteria slightly. It turns out that the Fast-2AFC task with the White&Small stimulus condition was particularly challenging for most observers, with several reporting that this condition was only a momentary flash that made perceptual judgements difficult. One observer (P5) exceeded the exclusion criterion of bandwidth σ>0.78 for this condition (σ=0.85). However, this larger bandwidth was consistent with the values seen in the rest of the sample, and this observer’s other results accorded well with the overall data. We therefore decided to retain this observer in the study. A control analysis using a stringent exclusion criterion (σ>0.5) confirmed that this did not substantially alter the conclusions (Supplementary Figures S1C and S2C).

Finally, we were instructed by the journal editor to collect additional data to bring the total sample size in Experiment 1 to N=24. Given that the project entailed several robustness analyses and generating figures, we adapted our analysis to automate the calculation of correlation Bayes Factors using custom MATLAB code, instead of using JASP.

## Results

Across two experiments, we aimed to better understand the discrepancies between previous studies in terms of the factors governing visual contextual size illusions. We quantified individual differences in illusion magnitudes and tested the effects of task and stimulus parameters on the Ebbinghaus illusion.

### Illusions with disc targets are related

Our first experiment used a battery of tasks to compare the magnitudes of our range of illusions. Averaged across observers, the illusion magnitudes varied considerably, with by far the strongest effect for the Mueller-Lyer and the weakest for the Ponzo illusion (Figure 1G), with the Ebbinghaus and Delboeuf magnitude at similar intermediate levels. We also quantified the mean bandwidths of psychometric curves as a reflection of objective discrimination thresholds. The bandwidth was largest (i.e., worst) for the Delboeuf and slightly elevated for the Ebbinghaus, but remained at similar levels for the other illusions and the objective discrimination tasks (Figure 1G).

Next, we calculated the pairwise correlations between the different illusion magnitudes and the bandwidths for objective disc and line discrimination (Figure 1H, see also Supplementary Figure S1A for scatter plots of each correlation). This confirmed the hypothesised primary correlation (I_1_) between the Ebbinghaus and Delboeuf illusion (r=0.50, BF_+0_=9.235). Surprisingly, the Ponzo illusion further correlated positively with the Delboeuf illusion (Delboeuf: r=0.55, BF_+0_=20.771). Additionally, we found a positive correlation between the Ponzo and the Ebbinghaus illusion, although statistical evidence for this association was inconclusive (r=0.29, BF_+0_=1.147). These findings contradict earlier findings by ourselves and others (Jastrzębowska et al., 2023; Schwarzkopf et al., 2011), but are in line with other recent studies (Mazuz et al., 2023). This at least partially rejects our preregistered hypothesis I_2_.

Delboeuf illusion magnitude also correlated strongly with objective thresholds (bandwidths) for discriminating disc stimuli (r=0.52, BF_+0_=11.840). We also observed positive correlations between disc discrimination and both the Ebbinghaus (r=0.41, BF_+0_=3.09) and Ponzo illusions (r=0.56, BF_+0_=22.471), the other illusions using disc targets. This therefore fully supports our hypothesis S_1_ about disc discrimination abilities and rejects hypothesis S_2_ (Table 1). Interestingly, we found moderate evidence for the null hypothesis of no relationship between thresholds for line discrimination and the Ebbinghaus illusion (r=-0.01, BF_+0_=0.240). For the other illusions, statistical evidence here was inconclusive (⅓<BF_+0_<3), except for the Ponzo where we found a positive correlation (r=0.44, BF_+0_=4.418). This means not even the Mueller-Lyer illusion correlated positively with this objective task, despite using the same line targets. This is evidence against our preregistered hypothesis M_1_, although these results remain inconclusive.

There was also evidence against any relationship between the Mueller-Lyer and the Ebbinghaus (r=-0.07, BF_+0_=0.203) and Delboeuf (r=-0.01, BF_+0_=0.248) illusions. The Mueller-Lyer also did not correlate with objective disc discrimination ability (r=0.06, BF_+0_=0.315). Evidence for other correlations involving Mueller-Lyer was inconclusive. As such, our results do not fully adjudicate between the two exploratory hypotheses M2 and M3 we specified for that illusion. While it was not linked with the assumed low-level effects (Ebbinghaus, Delboeuf) it remains unclear to what extent it is associated with the presumably higher-level Ponzo illusion. Finally, contrary to hypothesis S_3_, discrimination thresholds for lines and discs were effectively uncorrelated, although statistical evidence for the alternative or null hypothesis was inconclusive (r=0.31, BF_+0_=1.230).

Like many psychophysical studies on fundamental visual processing, our sample included the authors of this study. The pattern of results however remained qualitatively unchanged when restricting the analysis to only the naïve observers (Figure S1B). When applying a stricter exclusion criterion by removing observers whose discrimination bandwidth exceeded σ>0.5 in any condition, the main pattern of results also largely remains stable. However, only objective disc-width discrimination correlated with the Ponzo illusion in that analysis, and evidence for the null hypothesis generally became more conclusive in several comparisons that yielded inconclusive results in the main analysis. However, evidence for the associations between the Ponzo and, respectively, the Delboeuf illusions and line-width discrimination were inconclusive in this analysis (Figure S1C).

### Task modulates illusion magnitude

We next ran a battery of Ebbinghaus illusion measurements, using an Adjustment task, an Unconstrained-2AFC task (unlimited stimulus duration and free eye gaze), and a Fast-2AFC task (brief stimulus duration and instruction to maintain fixation). In this experiment, we also compared two different stimulus designs for the illusion (Figure 2A-B). Averaged across observers, illusion magnitudes varied substantially between the three tasks (Figure 2C; effect of task: BF_incl_=1235.192), confirming our preregistered hypothesis H_1_. Specifically, illusion magnitudes were lower when measured with the Adjustment task than with either the Unconstrained-2AFC (BF_10_=298163.190) or the Fast-2AFC task (BF_10_=2248.053). In contrast, average illusion magnitude did not differ between the two 2AFC tasks (BF_10_=0.295). These findings confirm hypothesis H_1A_ but reject hypothesis H_1B_.

To further explore these data, we also compared discrimination bandwidths for the 2AFC tasks (Figure 2D). Bandwidths were consistently smaller for the Unconstrained-2AFC than for the Fast-2AFC task (BF_10_=97071.983), meaning the latter was more difficult.

### Different stimulus parameters do not affect illusion magnitude

The stimuli with White&Small discs yielded the weaker illusions (BF_incl_=12.215), but this effect depended on the task as evidenced by an interaction between task and stimulus (BF_incl_=29.927). Specifically, while there was strong evidence for this difference in the Unconstrained-2AFC task (BF_10_=333.991), evidence was inconclusive in the Adjustment task (BF_10_=2.088), while in the Fast-2AFC task, there was compelling evidence against any difference between stimulus conditions (BF_10_=0.255). These findings broadly support our preregistered hypothesis H_3_, but only within the constraint that this depends on the task used to measure illusions. Importantly, this applies to the relative illusion effect, expressed as the ratio between test and reference stimulus diameters on a logarithmic scale. In terms of absolute size differences in degrees of visual angle, the illusion magnitude for the White&Small stimuli would be approximately half that of the Grey&Large stimuli.

The lack of any difference in illusion magnitude stands in contrast to a pronounced difference in discrimination ability. Bandwidths were consistently larger (worse) for White&Small than for Grey&Large discs (Figure 2D; BF_10_=283.791). Especially for the Fast-2AFC task, judgements became difficult for White&Small discs, to the extent that we decided to deviate from our preregistered exclusion criteria: the discrimination bandwidth for several observers exceeded our predefined criterion for this condition. However, we decided to include them, as this pattern was consistent with the whole sample and their other data were of high quality.

Illusion magnitudes for the two stimulus conditions were also strongly correlated, both averaged across tasks (r=0.85, BF_+0_=122344.050) as well as separately for the Adjustment (r=0.66, BF_+0_=69.598), Unconstrained-2AFC (r=0.86, BF_+0_=112975.446), and the Fast-2AFC task (r=0.76, BF_+0_=1083.176). Exploring this further, we found that the strength of correlations between stimulus conditions did not differ between tasks (all 95% confidence intervals overlapped zero).

### Brief stimulation and fixation affect illusion measurements

We then calculated the correlations between the illusion magnitudes across different tasks and stimulus conditions. This revealed that the Adjustment and Unconstrained-2AFC tasks were strongly correlated with one another (Figure 2E; all BF_+0_>33.652). In contrast, while the Fast-2AFC task also correlated positively with the Adjustment task, these correlations were significantly attenuated for the White&Small stimulus condition (δT=-2.93, 95^th^ percentile: - 0.93). For Grey&Large stimuli, we found the same trend, although this did not reach statistical significance (δT=-0.96, 95^th^ percentile: 0.44). These findings at least broadly support our preregistered hypotheses H_2_ about the individual differences between tasks. We observed the weakest correlation between the Fast-2AFC task with White&Small stimuli and the other conditions. While positive, evidence tended to favour supporting the presence of correlations (all BF_+0_>1), only the correlations for Grey&Large stimuli (with Adjustment: r=0.50, BF_+0_=7.680; with Unconstrained-2AFC: r=0.50, BF_+0_=8.498) and one for White&Small stimuli (with Unconstrained-2AFC: r=0.48, BF_+0_=6.279) conclusively supported this relationship with the other tasks. Scatter plots for these correlations are shown in Supplementary Figure S2A. Moreover, to rule out any undue bias due to inclusion of non-naïve (author) observers, we repeated this analysis restricted to naïve observers. This yielded qualitatively similar results (Supplementary Figure S2B). We also ran a robustness analysis with our stricter exclusion criterion by removing observers for whom discrimination bandwidth exceeded σ>0.5. Given the difficulty of the Fast-2AFC task with White&Small stimuli, this reduced the sample size considerably (N=14), resulting in fewer conclusive results; nevertheless, the general pattern of results was preserved (Supplementary Figure S2C).

### Delboeuf illusion shares large proportion of its variance with Ebbinghaus

We next estimated the test-retest reliability of illusion measurements from split-half data (Supplementary Figure S3). This showed that the measurements for most tasks and stimulus conditions were highly reliable. The Unconstrained-2AFC task, regardless of stimulus type, was most reliable, almost reaching ceiling (R_rel_=0.98). Interestingly, we found the lowest reliability for White&Small stimuli in the Adjustment (R_rel_=0.84) and the Fast-2AFC tasks (R_rel_=0.85). The reduced reliability in the Fast-2AFC task with White&Small stimuli was presumably due to poorer overall performance for this condition. However, the Fast-2AFC measurements for the Grey&Large stimuli were extremely reliable (R_rel_=0.93), even though this task was more challenging than both the Adjustment and Unconstrained-2AFC (Figure 2D). We further compared these reliability estimates statistically. This revealed that reliability for the Adjustment was significantly lower than for the Unconstrained-2AFC task (Grey&Large: δ_rel_= - 0.06, 95^th^ percentile: -0.01, White&Small: δ_rel_=-0.14, 95^th^ percentile: -0.06). However, while the Adjustment also trended to be less reliable than the Fast-2AFC condition, this difference was not significant (Grey&Large: δ_rel_=-0.01, 95^th^ percentile: 0.12, White&Small: δ_rel_=-0.01, 95^th^ percentile: 0.18).

It is worth noting that our Adjustment condition was based on the average of 10 trials for each stimulus conditions. This is considerably more data than is used in many similar studies in the literature, which have used only two trials per condition (e.g., Cretenoud et al., 2021; Grzeczkowski et al., 2017; Jastrzębowska et al., 2023). Averaging across fewer trials increases the variance and thus reduce internal reliability. To estimate the impact this choice in experimental parameters could have, we recomputed the test-retest reliability using only the first two trials for each observer. As expected, the reliability was lower than for the full dataset (Grey&Large: R_rel_=0.77, White&Small: R_rel_=0.66). Importantly, it is also lower than all 2AFC conditions (all δ_rel_≤-0.16) and this reduction was significant for all comparisons (all 95^th^ percentiles < 0), except for the Fast-2AFC task with White&Small stimuli (95^th^ percentile: 0.12).

Since the Fast-2AFC condition with Grey&Large stimuli was effectively identical to the Ebbinghaus task in Experiment 1 and used the same observers (except for one observer excluded based on preregistered criteria in the Experiment 2), we were able to obtain a separate reliability estimate. Comparing these measurements also revealed a strong correlation (albeit weaker than the internal reliability in Experiment 2) between illusion magnitudes (Figure 2F, black curve; r=0.82, BF_+0_=20931.952) but a substantially weaker correlation between discrimination bandwidths (Figure 2F, blue curve; r=0.43, BF_+0_= 3.474).

## Discussion

Our findings demonstrate a strong correlation between the magnitudes of the Ebbinghaus and Delboeuf illusions when using a classic two-alternative forced-choice (2AFC) procedure, as we used in our earlier work investigating the links between perception and neural processing (Schwarzkopf & Rees, 2013). These findings support previous suggestions that the Ebbinghaus and Delboeuf are essentially manifestations of the same underlying effect (Roberts et al., 2005; Sherman & Chouinard, 2016; Todorović & Jovanović, 2018; Urale & Schwarzkopf, 2023). Even though Ebbinghaus stimuli are considerably more complex - with multiple contextual stimuli defined by numerous parameters that likely interact with one another as well as with the target – the Delboeuf illusion magnitude still remarkably explains a significant amount of the variance in the Ebbinghaus.

This finding, however, contrasts with recent reports suggesting that these two illusions are not substantially related (Jastrzębowska et al., 2023). From our present findings, we deduce that this discrepancy could be due to the differences in methodology: previous research mostly used adjustment procedures. This approach objectively fails to control stimulus duration and retinotopic location. Potentially, it could also be skewed by cognitive and/or decision-making biases, given the extended response time to make a perceptual judgement (Morgan et al., 2013). Even in the few cases where previous studies employed 2AFC procedures, stimulus characteristics were not well controlled and were therefore also likely susceptible to decision confounds. Interestingly, the correlation between tasks in that earlier study (Cretenoud et al., 2021) was comparable to the correlations we found between Adjustment and Unconstrained-2AFC tasks in our second experiment. Importantly, the correlations between Adjustment and Fast-2AFC tasks were substantially lower. This discrepancy could call the reliability of these earlier studies into question. We will return to this point below.

Importantly, our present study took care to match the stimulus conditions between the illusions and discrimination tasks. We note that previous research taking a similar approach to closely match stimulus conditions also found strongly correlated estimates for the Ebbinghaus and Delboeuf illusions, even when using an adjustment task (Sherman & Chouinard, 2016). Matching stimuli is essential for drawing meaningful conclusions about how these perceptual effects relate to neural representations in visual cortex and how they may depend on local cortical magnification. Importantly, this also maximises the chance of observing any correlation between the illusions; all else being equal, the only remaining difference is the type of illusion used. Assuming these effects depend on visual cortex representations, any detectable relationship between these effects should be weakened if the conditions differ in terms of geometric dimensions and where in the visual field the stimuli are presented.

Therefore, it is perhaps not surprising that we also found correlations between the Ponzo illusion and the other two using disc targets (Delboeuf and Ebbinghaus, although for the latter results were not statistically conclusive). What is more, all these illusions correlated positively with the objective ability to discriminate disc widths. While there have been reports of an association between the Ebbinghaus and Ponzo (Grzeczkowski et al., 2017; Mazuz et al., 2023), other research has found weak association between these illusions at best (Jastrzębowska et al., 2023; Schwarzkopf et al., 2011), consistent with the idea that the Ebbinghaus is largely mediated by local interactions between V1 neurons (Jaeger et al., 2014; Jaeger & Grasso, 1993; Jaeger & Klahs, 2015; Roberts et al., 2005; Schwarzkopf & Rees, 2013; Song et al., 2011; Todorović & Jovanović, 2018; Urale & Schwarzkopf, 2023), while the Ponzo involves interpretations of the three-dimensional scene – a process that highly likely recruits higher-order processes (Chen et al., 2018; Song et al., 2011). Our findings now add further nuance to this interpretation. As with most illusions, the Ponzo likely represents a mixture of both low- and high-level processes. The correlation reported here captures the former variance, which is shared with the Ebbinghaus and Delboeuf, while unexplained variance still remains that could stem from higher-order mechanisms.

Given our matched stimulus characteristics, the most striking aspect of our results is in fact the *absence* of positive correlations, supported by statistical evidence for the null hypothesis, most notably when comparing the Mueller-Lyer illusion with the others. The neural mechanisms mediating the Mueller-Lyer remain unclear, as it could involve both high-level interpretation of three-dimensional perspective (Gregory, 1966) and low-level image filtering (Carrasco et al., 1986; Ho & Schwarzkopf, 2022; Morgan & Glennerster, 1991). Our results suggest that if the Ponzo and the Mueller-Lyer both involve higher-level processes, these are unlikely to be mediated by the same mechanism. Curiously, the Mueller-Lyer also did not correlate with the objective line-length discrimination, and the two objective discrimination tasks also appear unrelated (although the result for line-length discrimination was statistically inconclusive). This indicates that the perceptual judgement underlying the Mueller-Lyer is rather different from that mediating the other tasks. The Mueller-Lyer is also by far the strongest and most consistent of the illusions we tested. In future work, it could be interesting to design a Mueller-Lyer illusion with disc targets (e.g., surrounded by a crown of arrowheads) to ascertain if the absence of correlation observed here is due to the different target stimuli.

In our second experiment, we compared Ebbinghaus illusion measurements between different tasks. We found that estimates from Adjustment tasks were very similar to those obtained with the Unconstrained-2AFC task, where observers could inspect stimuli for as long as they needed and were free to move their eyes. In contrast, estimates were far less consistent between these tasks and the Fast-2AFC task, where eye gaze and stimulus duration were tightly controlled. It is tempting to argue that these diminished correlations are a trivial consequence of poorer discrimination performance for the Fast-2AFC task. Indeed, discrimination thresholds for this task, especially with the stimuli comprising White&Small discs, were consistently higher (worse). However, Ebbinghaus illusion measurements, at least for Grey&Large stimuli, were highly reliable and revealed strong correlations with the Delboeuf and Ponzo illusions in Experiment 1. This again stands in contrast to a previous study using adjustment tasks, which failed to find any compelling associations between these illusions (Jastrzębowska et al., 2023). We also noted high correlations between the two stimulus designs for the Fast-2AFC task. Therefore, we surmise that the Fast-2AFC task can yield very robust measurements. The lower correlation between the Fast-2AFC task and Adjustment estimates, compared to the Unconstrained-2AFC task, likely results from the latter’s shared critical characteristics with Adjustment: Unlimited stimulus presentation and unconstrained eye movements. As laid out in the Introduction, this could imply cognitive factors also play a greater role: By giving observers long time windows to consider the stimuli, both Adjustment and Unconstrained-2AFC arguably involve more cognition than a speeded forced-choice decision.

On the other hand, it could also be argued that the Fast-2AFC task entails a memory component because the stimuli have disappeared by the time the observer can respond, potentially involving an additional factor not involved in the unconstrained tasks. Because the stimuli vanish before the observer can respond, observers must rely on an internal representation of the stimuli to make a perceptual judgement. This could involve iconic or working memory, although with brief stimuli and speeded responses the former is perhaps more likely. Judging which of two simultaneous stimuli is larger is not all that different from judging the orientation of a briefly flashed grating – yet the latter task is rarely regarded as a test of memory function. Nevertheless, one could consider this a higher-level process than reproducing a perceptual match in an adjustment task. On balance, however, the long decision window to inspect stimuli and make an adjustment still allows more time for cognition to play a role than speeded decisions in a forced-choice task. And even if there were other high-level mechanisms at play in forced-choice tasks, as we already discussed, briefly presented stimuli inevitably provide better control over eye fixation and stimulus properties.

In this context, we note that our Adjustment data, despite being based on an unusually large number of trials for such experiments, were still less reliable than most 2AFC measurements (except for the difficult Fast-2AFC White&Small condition). This was particularly the case when using only the first two adjustment trials to estimate illusion magnitude, similar to how this was done in previous studies (e.g., Cretenoud et al., 2021; Grzeczkowski et al., 2017; Jastrzębowska et al., 2023). Low internal reliability for one variable diminishes the sensitivity for detecting associations with other variables. This is particularly problematic when testing associations between psychophysical measures and highly variable neurophysiological variables. The lower reliability of the Adjustment tasks is also a further indication of differing contributing processes that underlie the illusion measurement using different tasks.

Indeed, by fixing eye position and limiting stimulus durations the Fast-2AFC design likely engages different neuronal populations in visual cortex than the Adjustment and Unconstrained-2AFC tasks. Only the Fast-2AFC task allows for relatively tight control over this aspect. It is difficult to ascertain which regions of the visual field – and therefore which corresponding neuronal populations – are involved during perceptual judgements in the Adjustment and Unconstrained-2AFC tasks. Whenever an observer foveates one of the two target stimuli, it is represented with the highest spatial resolution in foveal cortex, while the other stimulus is relegated to a peripheral location with reduced resolution. Consequently, the eccentricity of this fellow stimulus would then be double that of both, the test and reference stimuli, in the Fast-2AFC task. If the cortical representation matters to the perceptual effect, then the illusion estimate must inevitably differ. Notably, stimuli are typically perceived as smaller in peripheral vision (Anstis, 1998; Moutsiana et al., 2016; Newsome, 1972; von Helmholtz, 1924). By directing fixation to the midpoint between the two stimuli, the Fast-2AFC controls for this aspect, while the illusion estimates in the other two tasks may include this general perceptual bias. The situation is further complicated by the fact that observers can alternately foveate the two targets or occasionally fixate between them. It is difficult to determine which of these states contribute most to the illusion estimate. Either way, it also seems likely that these unconstrained tasks leave more room for decision factors to influence the response. While not directly comparable, a study comparing the vertical-horizontal (bisection) illusion, reported a profound effect of stable eye fixation (Chouinard et al., 2017).

In that light, it is puzzling that several previous studies using adjustment tasks found so few correlations between different illusions (Axelrod et al., 2017; Grzeczkowski et al., 2017; Jastrzębowska et al., 2023). If these tasks were dominated by decision factors, should they not in fact show greater consistency with each other? For example, if these results reflected the conservativism of their decision criterion to reproduce a perceptual match, why would this criterion differ so wildly between these illusions? Such a decision criterion could be considered a common factor to these illusions, possibly related to personality factors (Makowski et al., 2023) – the accumulated data across the literature speak for themselves that such a factor either does not exist or only explains such a small amount of variance that could not be reliably detected even in many large-scale studies.

Certainly, such a decision factor should not be reflected by the functional architecture in V1. Rather, local properties of V1, which vary with visual field location and geometric stimulus dimensions, should exert more specific effects on perception. This does not mean illusion estimates do not generalise up to a point across stimulus characteristics and visual field locations – some generalisation is expected, as neural properties also generalise. For instance, the cortical magnification factor or population receptive field sizes at 3° eccentricity must at least correlate somewhat with those at 1° eccentricity. In our study, the relative illusion estimates for the two stimulus conditions in the second experiment were indeed very similar, despite substantial differences in absolute change in degrees of visual angle. More generally, although task-related differences were significant, illusion estimates remained relatively well-correlated across tasks. Up to a point, it is feasible to use comparable stimuli and experimental parameters in perceptual studies and explore how the effects relate to the brain. However, the more differences are introduced into the design, the greater the variability becomes and the less likely it is that two measures correlate.

### Limitations

The sample size in our original study (Experiment 1: N=12, Experiment 2: N=11) was small for studying individual differences. Our preregistered sampling plan prescribed stopping data collection when we obtained strong evidence either for (BF_+0_>10) or against (BF_+0_<0.1) a correlation between the Ebbinghaus and Delboeuf illusion magnitude. Given similar previous research (Jastrzębowska et al., 2023; Sherman & Chouinard, 2016), we anticipated that we would require a larger sample, setting our maximum at N=90. However, we already obtained compelling evidence that these two illusions are correlated after the second testing milestone. As explained in the Introduction, we deliberately designed our illusion stimuli to closely match fundamental attributes between conditions, thereby controlling for stimulus parameters known to affect these illusions (Jaeger et al., 2014; Jaeger & Grasso, 1993; Jaeger & Klahs, 2015; Roberts et al., 2005; Todorović & Jovanović, 2018; Urale & Schwarzkopf, 2023). This could have contributed to high similarity in these illusion estimates. Either way, our sample size was evidently sufficient to obtain conclusive results. Nevertheless, we addressed this potential limitation by collecting data from 12 additional observers, effectively doubling the sample size. While the study can therefore not be considered purely confirmatory, this ensured more conclusive results for many of the hypotheses we tested.

It could also be argued that our sample was demographically too homogenous, especially considering the small number of observers, leading to potential biases. As is common in psychophysics studies, the three non-naïve authors also participated in the experiment. Our control analyses confirmed that their inclusion did not qualitatively affect the results. While our observers were ethnically comparably diverse and their ages ranged from young adults to middle-aged, the majority were female. Most observers were also either present or recent students/staff at the University of Auckland. The Ebbinghaus illusion reportedly varies with age, with weaker illusions in children (Doherty et al., 2010; Weintraub, 1979) and older age compared to young adults (Mazuz et al., 2024). It reportedly also varies across cultural contexts (Bremner et al., 2016; de Fockert et al., 2007). Thus, it is certainly possible that our limited sample restricted the range of illusion magnitudes one would find in a broad global sample.

However, this would also be accompanied by additional confounds. A broader age range, for example, could introduce variance related to changes in general visual abilities, while a global population likely varies in terms of their practice, concentration, and skill in performing such computerised tasks. This would inevitably reduce the reliability of these measures and weaken the strength of associations between illusions. In turn, this would be contrary to the aims of our study, which sought to isolate the existence of shared factors in *low-level visual processing* under well-controlled conditions. Just as we designed our stimuli to be closely matched, our comparably homogenous sample enabled us to reveal the role of stimulus parameters and task design without contamination from other confounding factors like age, gender, cultural context, or visual diet. This is a prerequisite step before larger global studies that can later establish which additional factors contribute to illusion estimates.

Another potential limitation of our experiment is that the task instructions for the 2AFC tasks inadvertently differed from those in the Adjustment tasks. In the former, observers were instructed to choose the target disc that “was larger.” In the latter, the instruction was to adjust the test disc size until both discs “appeared equal.” It is possible that the exact instructions can affect how observers respond, especially when studying neurodivergent observers. However, we consider it unlikely that this played a significant role in our experiments. Illusion estimates for the Adjustment and Unconstrained-2AFC tasks were strongly correlated despite this discrepancy in task instructions. By contrast, correlations between these two tasks and the Fast-2AFC were considerably weaker, even though the Fast-2AFC task with Grey&Large stimuli was highly reliable, and produced consistent measurements irrespective of the stimulus design. This suggests that this task probed separate processes not tapped by the other two tasks.

### Conclusions

Any illusion estimate must reflect multiple separate mechanisms (some of which probably also interact), ranging from basic perceptual processing to complex decision-making. Therefore, any notion of a single common factor that could be shared by all illusions, even all spatial contextual illusions of size as studied here, seems implausible. Even for potential common factors like general intelligence or personality traits, extraneous variables can influence the outcome of specific tasks in individuals – for example, a colour-blind person would find it difficult to pick a correct target defined by colours even if their intelligence is not a limiting factor. More importantly, a hypothetical common factor, equivalent to general intelligence, that determines a person’s susceptibility to visual illusions is based on circular logic: To recognise that their percept disagrees with the physical reality of the stimulus, an observer must have the ability to separate the perceived stimulus from the retinal image. It seems improbable that such a general process would have evolved, if these illusions reflect stimulus processing necessary for functional vision. Our original hypothesis linking the Ebbinghaus, Delboeuf, and similar contextual effects like the tilt illusion, to cortical distance, is based on the idea that they result from feedforward filtering of the visual input by receptive fields (Mareschal et al., 2010; Moutsiana et al., 2016; Schwarzkopf et al., 2011; Schwarzkopf & Rees, 2013; Song et al., 2015; Song, Schwarzkopf, & Rees, 2013; Urale & Schwarzkopf, 2023). This suggests that the information about the true stimulus has been filtered out at higher stages of processing. Any measure that reflects this neural processing of visual input, such as measuring cortical magnification and/or population receptive field size, should therefore explain some of the variance in these specific illusions – but not all of it, and perhaps not even the majority.

However, this also means that the opposite implication suggested by some studies (Cretenoud et al., 2021; Grzeczkowski et al., 2017; Jastrzębowska et al., 2023) is even less plausible, namely that there are *no* factors shared at least by some illusions. Taken to its logical conclusion, this notion would imply that each illusion is mediated by entirely separate neural mechanisms. This is akin to the concept of a “grandmother neuron” in memory or object recognition. Just as it seems unlikely that there is only one cell (or a small population of cells) that encodes a specific person’s identity, it seems highly implausible that there are wholly separate neural mechanisms for the Ebbinghaus, Delboeuf, Ponzo, or Mueller-Lyer illusions. An alternative explanation put forth for the claimed lack of correlation between different illusions is that the barrier to understanding their neural basis lies in the “computational complexity” (Ozkirli et al., 2025). However, this merely restates the point we made already that multiple interacting processes are involved in *any* behaviour, including measuring illusions. As such, it is likely that some related effects only share a small amount of variance. Revealing such relationships is however not impossible – rather, it requires high statistical power, achieved through increasing sample sizes *and* using the most robust methods for measuring the illusion.

Many visual illusions are discovered by accident or deliberately designed based on prior knowledge of how visual processing is thought to work. They tap into underlying mechanisms – and thus, illusions that partially engage the same mechanisms must also be statistically related to each other. Characterising the specific parameters governing the Ebbinghaus and Delboeuf illusions has led researchers to propose that these are the same perceptual process (Roberts et al., 2005). One factor determining this effect may be cortical magnification (Urale & Schwarzkopf, 2023), as originally suggested by individual differences (Schwarzkopf & Rees, 2013). However, this does not mean that no other factors are involved – more research is necessary to identify these factors.

For example, it remains possible that the Ebbinghaus illusion – despite being profoundly determined by basic stimulus attributes (Jaeger et al., 2014; Jaeger & Grasso, 1993; Jaeger & Klahs, 2015; Roberts et al., 2005; Sherman & Chouinard, 2016; Todorović & Jovanović, 2018; Urale & Schwarzkopf, 2023) – also includes a genuine size contrast factor. This could underlie at least some similarity effects reported for this illusion (Choplin & Medin, 1999; Coren & Enns, 1993; Deni & Brigner, 1997). The contribution of component mechanisms could also vary between individuals and with demographic factors. While the Ebbinghaus varies between populations (Bremner et al., 2016; de Fockert et al., 2007) and in children (Doherty et al., 2010), such differences could result from changes in the size contrast factor or decision-making processes (Weintraub, 1979), rather than being explained by changes in cortical magnification. In fact, at least the macroscopic and mesoscopic functional architecture of early visual cortex is close to adult levels already at a young age (Dekker et al., 2019), although more subtle developmental changes continue to occur over the course of development (Himmelberg et al., 2023). This underscores the importance of experimental paradigms that can disentangle these factors.

Classical or novel forced-choice procedures (e.g., Finlayson et al., 2017; Jogan & Stocker, 2014; Morgan et al., 2013; Patten & Clifford, 2015) with closely matched stimulus properties, as used in our study, are a prerequisite for testing these low-level geometric factors of subjective perceptual experience. However, this does not mean that procedures like adjustment cannot yield important insights. Indeed, for determining the role of separate factors, a combination of different methods (Manning et al., 2017) may be the most fruitful approach. Critically, researchers should use the methods most appropriate for testing the role of specific factors, aiming for theoretical coherence and statistical reliability. Moreover, when making claims about the replicability of previous illusion measurements (or the lack thereof), it is vital to use the same procedures to measure perception.

## Data availability

All stimulus presentation materials, data, and analysis code are publicly available: https://osf.io/5zs7y and https://osf.io/b3h4x

## Acknowledgements

We thank Adina Giurgiu and Arash Chaparian for their help in setting up our new testing room.

## Supplementary Information

**Figure S1.**
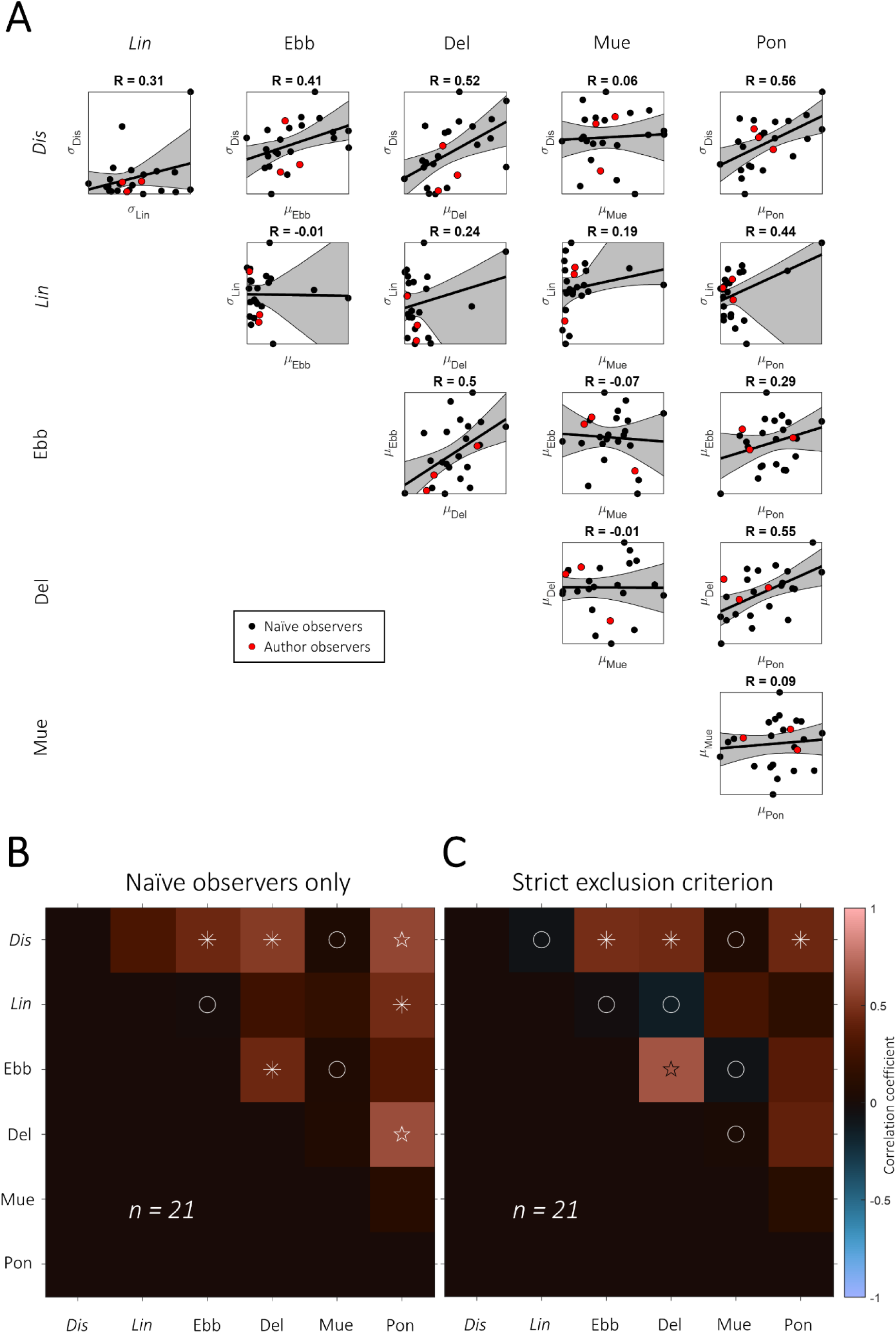
A. Scatter plots underlying the correlation matrix of data from Experiment 1 shown in Figure 1H. Each dot plots illusion magnitude and/or discrimination bandwidth for an individual observer from a given condition. Pearson correlation statistics are shown at the top of each panel (p-values rounded to three decimal points). Black line and shaded area denote linear regression and its 95% confidence interval. Black dots: Naïve observers. Red dots: Non-naïve (author) observers. B. Correlation matrix from Experiment 1 but excluding non-naïve (author) observers. C. Correlation matrix from Experiment 1 but excluding observers whose bandwidth (σ) exceeded 0.5 in any condition. Correlations compared the magnitudes for the four illusions, and discrimination abilities. All conventions as in Figure 1H. Ebb: Ebbinghaus. Del: Delboeuf. Mue: Mueller-Lyer. Pon: Ponzo. Dis: disc width. Lin: line length.

**Figure S2.**
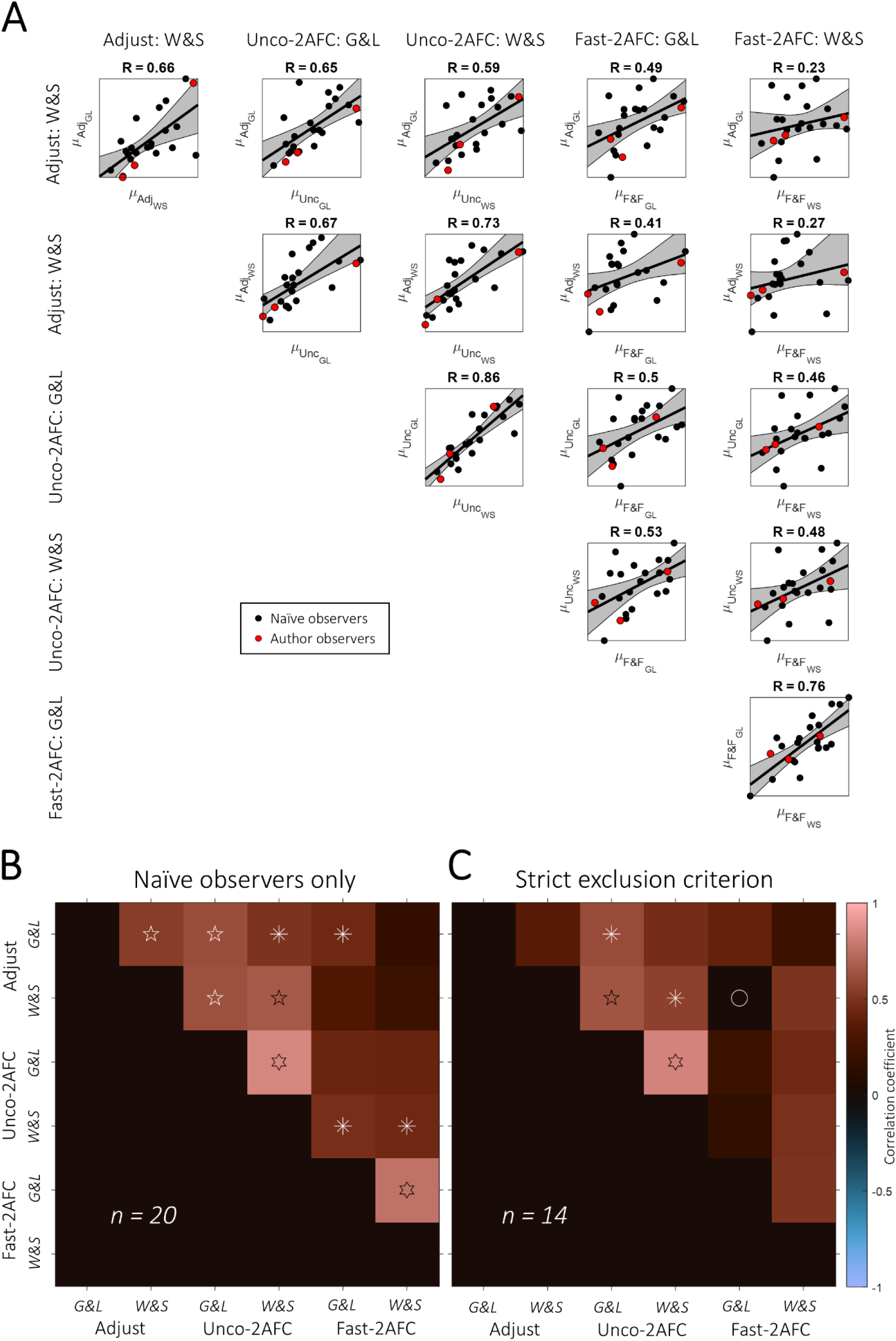
A. Scatter plots underlying the correlation matrix of data from Experiment 2 shown in Figure 2E. Each dot plots illusion magnitude for an individual observer from a given task and stimulus condition. Black line and shaded area denote linear regression and its 95% confidence interval. Pearson correlation statistics are shown at the top of each panel (p-values rounded to three decimal points). Black dots: Naïve observers. Red dots: Non-naïve (author) observers. Adj: Adjustment task. Unc: Unconstrained-2AFC task. F&F: Fast-2AFC task with fixation. B. Correlation matrix from Experiment 2 but excluding non-naïve (author) observers. C. Correlation matrix from Experiment 2 but excluding observers whose bandwidth (σ) exceeded 0.5 in any condition. All conventions as in Figure 2E.

**Figure S3.**
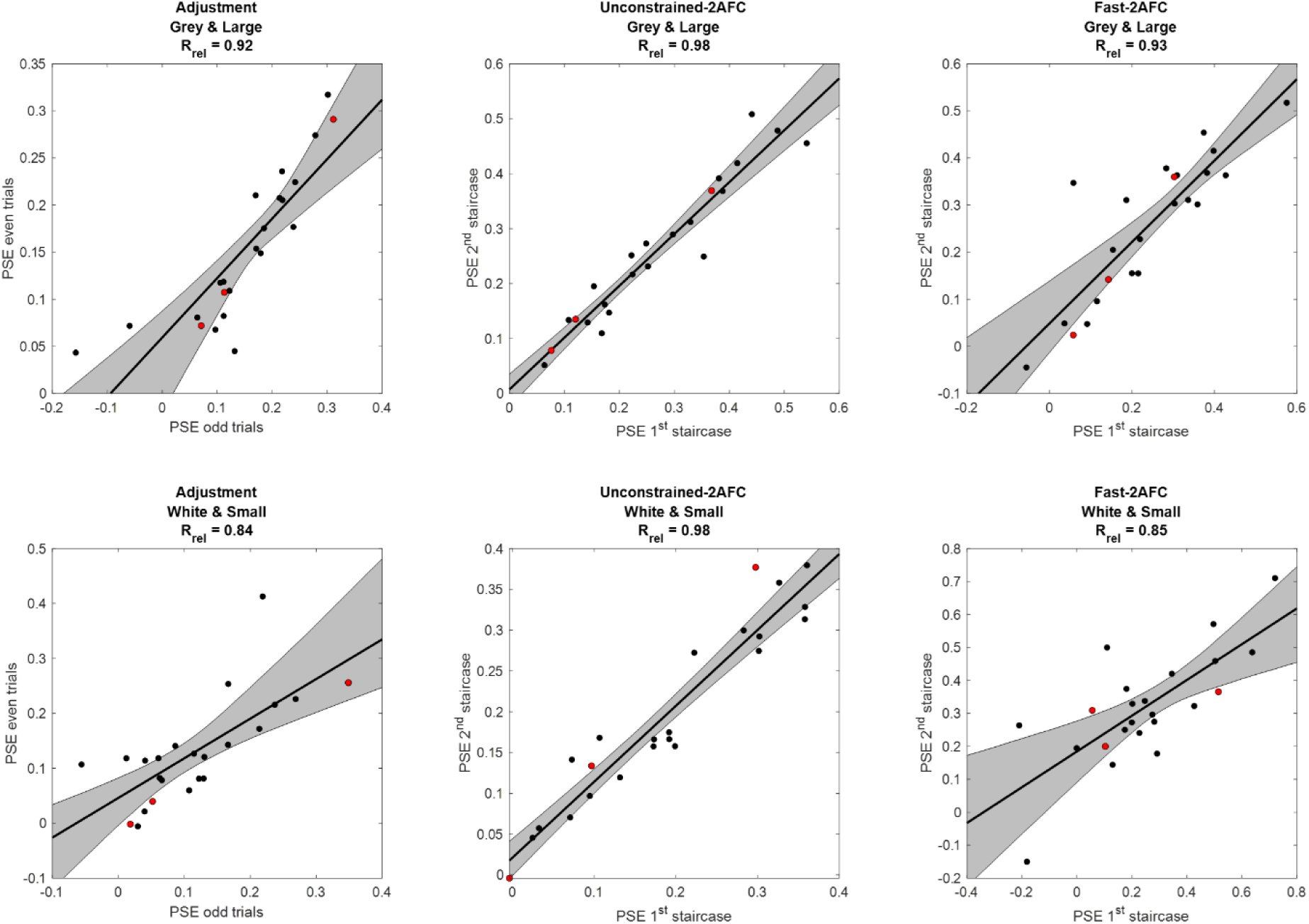
Scatter plots of the test-retest reliability for illusion magnitudes in each task and stimulus condition from Experiment 2. Each black dots denotes the illusion magnitude (point of subjective equality, PSE) estimated for a given observer from the second half of trials against the first half. In the Adjustment tasks, data were split into odd and even numbered trials. In the 2AFC tasks, separate psychometric curves were fit to the two independent, interleaved staircases. R_rel_ shows the test-retest correlation extrapolated for the full data set using Spearman-Brown prophecy formula (see Materials and Methods). Black line and shaded area denote linear regression and its 95% confidence interval.

